## Supplementary data for "Activated regulatory T-cells, dysfunctional and senescent T-cells dominate the microenvironment of pancreatic cancer"

### Tables

*Table 1: Patient data.*
PanIn= Pancreatic intraepithelial neoplasm. IPMN= Intraductal papillary mucinous neoplasm.

| Patient | Sex | Age | TNM stage | Size of primary | Background pancreas related pathology | Stent before surgery | Other co-morbidities | Baseline ca19.9 |
|---|---|---|---|---|---|---|---|---|
| 1 | Male | 71 | pT3 N2 (11/52) L1 V1 R1(Posterior, Medial) | 45mm | PanIn | No | No | N/A |
| 2 | Male | 72 | pT2 N1 (3/36) L1 V1 R0 | 28mm | PanIn | Yes | Diabetes mellitus, Acute kidney injury (Stage 1), Ex-smoker. | N/A |
| 3 | Female | 51 | pT2 N2 (12/66) M0 L1 V1 R2(Medial) | 26mm | IPMN | No | Diabetes mellitus. | Ca19.9 - 18 IU/mL |
| 4 | Male | 80 | pT3 N0(0/22 + 0/1 = 0/23) M0 L0 V0 Pn1 R0 | 15mm | No | No | Cancer of prostate. | Ca19.9 - 1,356 U/mL |
| 5 | Female | 73 | R1 (medial x 2 & pancreatic RM) pT3 L1 V0 Pn1 N2 (16/37) | 42mm | No | Yes | No | N/A |
| 6 | Female | 67 | pT3, N2 (10/68 M0, L1 V1 R1 anterior medial) | 28mm | PanIn | No | Inactive: Ex-smoker | Ca19.9 - 147 U/mL |
| 7 | Male | 67 | R1 (posterior) pT2 L0 V1 Pn1 N1(1/19) | 24mm | PanIn | No | Smoker. | N/A |
| 8 | Male | 69 | pT2 N2 (9/18) M0 L1 V1 R0 | 28mm | PanIn | No | Diabetes mellitus, Ex-smoker. | N/A |

*Table 2: CYTOF panel*

| Antigen | Metal Tag | Clone | Vendor, catalogue no. |
|---|---|---|---|
| CD45 | 89 Y | HI30 | Fluidigm, 3089003B |
| CD14 Qdot | Qdot655 (114 Cd) | TuK4 | ThermoFisher, Q10056 |
| EpCAM | 141 Pr | 9C4 | Fluidigm, 3141006B |
| CD19 | 142 Nd | HIB19 | Fluidigm, 3142001B |
| CD127 | 143 Nd | A019D5 | Fluidigm, 3143012B |
| HLA-A,B,C | 144 Nd | W6/32 | Fluidigm, 3144017B |
| CD4 | 145 Nd | RPA-T4 | Fluidigm, 3145001B |
| CD8a | 146 Nd | RPA-T8 | Fluidigm, 3146001B |
| CD39 | 147 Sm | A1 | Biolegend, 328221 |
| ICOS | 148 Nd | C398.4A | Fluidigm, 3148019B |
| CD25 | 149 Sm | 2A3 | Fluidigm, 3149010B |
| OX40 | 150 Nd | ACT35 | Fluidigm, 3150023B |
| CD103 | 151 Eu | BerACT8 | Fluidigm, 3151011B |
| CD66b | 152 Sm | 80H3 | Fluidigm, 3152011B |
| Tigit | 153 Eu | MBSA43 | Fluidigm, 3153019B |
| Tim-3 | 154 Sm | F382E2 | Fluidigm, 3154010B |
| CD27 | 155 Gd | L128 | Fluidigm, 3155001B |
| PD-L1 | 156 Gd | 29E.2A3 | Fluidigm, 3156026B |
| CD33 | 158 Gd | WM53 | Fluidigm, 3158001B |
| GITR | 159 Tb | 621 | Fluidigm, 3159020B |
| CD28 | 160 Gd | CD28.2 | Fluidigm, 3160003B |
| CTLA-4 | 161 Dy | 14D3 | Fluidigm, 3161004B |
| FoxP3 | 162 Dy | PCH101 | Fluidigm, 3162011A |
| CD56 | 163 Dy | NCAM16.2 | Fluidigm, 3163007B |
| CD15 | 164 Dy | W6D3 | Fluidigm, 3164001B |
| Lag-3 | 165 Ho | 11C3C65 | Fluidigm, 3165037B |
| CCR7 | 167 Er | G043H7 | Fluidigm, 3167009A |
| CD40L | 168 Er | 2431 | Fluidigm, 3168006B |
| CD45RA | 169 Tm | HI100 | Fluidigm, 3169008B |
| CD3 | 170 Er | UCHT1 | Fluidigm, 3170001B |
| Granzyme B | 171 Yb | GB11 | Fluidigm, 3171002B |
| Ki67 | 172 Yb | B56 | Fluidigm, 3172024B |
| 4-1BB | 173 Yb | 4B4-1 | Fluidigm, 3173015B |
| HLADR | 174 Yb | L243 | Fluidigm, 3174001B |
| PD-1 | 175 Lu | EH12.2H7 | Fluidigm, 3174020B |
| CD57 | 176 Yb | HCD57 | Fluidigm, 3176019B |
| CD16 | 209 Bi | 3G8 | Fluidigm, 3209003B |

**Supplementary Data**

**Supplemental Data File 1:** Top differentially expressed genes by $75^{th}$ percentile and means for Exhausted, Senescent and T-reg metaclusters.
**Supplemental Data File 2:** Differential expression analysis for Exhausted, Senescent and T-reg metaclusters.
**Supplemental Data File 3:** Reference pancreas and immune gene lists
**Supplemental Data File 4:** T-cell complete gene list

**Figure 1- figure supplement 1: Individual patients' immune populations**. Expression of clustering markers on top of a FlowSOM tree showing the relationship between metaclusters, for the pooled single cell data (top). Population frequency is presented in the pie chart in each circle along the tree, the background colour represents the metacluster identity. Metaclusters in these trees have higher granularity to allow for identifying additional features in the data compared to the summary in main figure the corresponding between here and main figure are: *MC10 (in Fig. 1D) = MC10+12 (in S1)*. (Bottom) FlowSOM trees showing the individual metacluster frequency for each patient. The size of the pie chart corresponds to the frequency relative frequency. (Expression profile: from low (Dark blue) to high (Deep red)).

**Figure 1- figure supplement 2. Individual patients' NK populations**. Expression of clustering markers on top of a FlowSOM tree showing the relationship between metacluster, for the pooled single-cell data (top; ~2,500 cells). Population frequency is presented in the pie chart in each circle along the tree, the background colour represents the metacluster identity. Metaclusters in these trees have higher granularity to allow for identifying additional features in the data compared to the summary in main figure the corresponding between here and main figure are: *MC2 (in Fig. 1E) = MC2+4 (in S2), MC6=MC7+8+9.* (Bottom) FlowSOM trees showing the individual metacluster frequency for each patient. The size of the pie chart corresponds to the frequency relative frequency. (Expression profile: from low (Dark blue) to high (Deep red)).

**Figure 1- figure supplement 3. Individual patients' Granulocyte populations.** Expression of clustering markers on top of a FlowSOM tree showing the relationship between metacluster, for the pooled single-cell data (top; ~36,000 cells). Population frequency is presented in the pie chart in each circle along the tree, the background colour represents the metacluster identity. Metaclusters in these trees have higher granularity to allow for identifying additional features in the data compared to the summary in main figure the corresponding between here and main figure are: *MC3 (in Fig. 1F) = MC3+4+6 (in S3).* (Bottom) FlowSOM trees showing the individual metacluster frequency for each patient. The size of the pie chart corresponds to the frequency relative frequency. (Expression profile: from low (Dark blue) to high (Deep red)).

**Figure 1- figure supplement 4. Individual patients' Mononuclear Phagocytes populations.** Expression of clustering markers on top of a FlowSOM tree showing the relationship between metacluster, for the pooled single-cell data (top; ~20,260 cells). Population frequency is presented in the pie chart in each circle along the tree, the background colour represents the metacluster identity. Metaclusters in these trees have higher granularity to allow for identifying additional features in the data compared to the summary in main figure the corresponding between here and main figure are: *MC2 (in Fig. 1G) = MC2+9 (in S4); MC3= MC3+4; MC4= MC5+6.* (Bottom) FlowSOM trees showing the individual metacluster frequency for each patient. The size of the pie chart corresponds to the frequency relative frequency. (Expression profile: from low (Dark blue) to high (Deep red)).

**Figure 1- figure supplement 5. Immune-complexes and MDSC observed in peripheral blood of PDAC patients.** (A) 124,000 CD45$^{+}$ cells pooled from 8 patients and viSNE analysis using main cell lineage markers was performed to identify the main immune cell populations. The main immune populations coloured and labelled by FlowSOM. Bar plots of metacluster frequencies in each patient. Inset shows the lower frequency metaclusters. Heatmap of FlowSOM metaclusters of CD45$^{+}$ cells; rows represent metaclusters from combined single cells across patients. CD66$^{+}$ represent low density neutrophils and CD14$^{+}$CD3$^{+}$ are T-cell/monocyte complexes. (B) ~12,750 NK cells were pooled and 13 different metaclusters identified with FlowSOM. Bar plots show metacluster frequencies. Inset shows the lower frequency metaclusters. Expression profile is presented in the heatmap. Most metacluster are Granzyme B$^{+}$, indicative of cytolytic capacity. (C) ~42,700 Mononuclear Phagocyte cells were pooled and 13 different metaclusters identified with FlowSOM. Bar plots show metacluster frequencies. Inset shows the lower frequency metaclusters. Expression profile is presented in the heatmap. The main metacluster appear to be MDSCs. All bar plots are median and the individual dots are individual patients. Heatmaps are normalised for each marker.

**Figure 2- figure supplement 1. Individual patients' CD8$^{+}$ T-cells populations.** Expression of clustering markers on top of a FlowSOM tree showing the relationship between metacluster, for the pooled single-cell data (top). Population frequency is presented in the pie chart in each circle along the tree, the background colour represents the metacluster identity. Metaclusters in these trees have higher granularity to allow for identifying additional features in the data compared to the summary in main figure the corresponding between here and main figure are: *MC1 (in Fig. 2A)= MC4+21+24+25+27+28+29 (in S6); MC2= MC5+20+23+26; MC3= MC18+MC19; MC4= MC2+3+14; MC6= MC12+13+17; MC9= MC7+8; MC10= MC6+MC11*. (Bottom) FlowSOM trees showing the individual metacluster frequency for each patient. The size of the pie chart corresponds to the frequency relative frequency. (Expression profile: from low (Dark blue) to high (Deep red)).

**Figure 2- figure supplement 2. Individual patients' CD4$^+$ T-cells populations.** Expression of clustering markers on top of a FlowSOM tree showing the relationship between metacluster, for the pooled single-cell data (top). Population frequency is presented in the pie chart in each circle along the tree, the background colour represents the metacluster identity. Metaclusters in these trees have higher granularity to allow for identifying additional features in the data compared to the summary in main figure the corresponding between here and main figure are: *MC1 (in Fig. 2B)= MC1+2+3+8+13 (in S7); MC3= MC11+12; MC4=MC10+14; MC8=MC4+5.* (Bottom) FlowSOM trees showing the individual metacluster frequency for each patient. The size of the pie chart corresponds to the frequency relative frequency. (Expression profile: from low (Dark blue) to high (Deep red)).

**Figure 2- figure supplement 3. Individual patients' Treg populations.** Expression of clustering markers on top of a FlowSOM tree showing the relationship between metacluster, for the pooled single-cell data (top). Population frequency is presented in the pie chart in each circle along the tree, the background colour represents the metacluster identity. *MC1 (in Fig. 2C)= MC14+15 (in S8); MC3= MC1+2+5; MC6=MC3+4+6+13; MC7=MC9+10.* (Bottom) FlowSOM trees showing the individual metacluster frequency for each patient. The size of the pie chart corresponds to the frequency relative frequency. (Expression profile: from low (Dark blue) to high (Deep red)).

**Figure 2- figure supplement 4. Similar T-cell signatures of senescence and suppression to tumours are observed in peripheral blood**. (A) ~18,270 $CD8^+$ T-cells were pooled and 13 metaclusters identified with FlowSOM and visualised on viSNE plot. Metaclusters' relative abundance is shown in the bar plot. Inset shows the lower frequency metaclusters. The heatmaps show the expression profile of immune checkpoints in the different metaclusters. Note that the major $CD8^+$ populations are senescent and central memory cells. (B) ~38,350 $CD4^+$ T-cells were pooled and 14 metaclusters identified with FlowSOM and visualised on viSNE plot. Metaclusters' relative abundance is shown in the bar plot. Inset shows the lower frequency metaclusters. The heatmaps show the expression profile of immune checkpoints in the different metaclusters. The major populations are effector and central memory cells, followed by regulatory ($Foxp3^+$) T-cells. Two patients also show a signature of senescent cells in the periphery. (C) ~3,600 $CD4^+$ regulatory T-cells were pooled and 10 metaclusters identified with FlowSOM and visualised on viSNE plot. Metaclusters' relative abundance is shown in the bar plot. Inset shows the lower frequency metaclusters. The heatmaps show the expression profile of immune checkpoints in the different metaclusters. There is a noticeable naïve metacluster, and the activated metacluster with similar charactertics to the tumour is observed at lower frequency. Bar plots are medians and each dot represents a patient. Heatmaps were normalised across all T-cell populations per marker. Hierarchal clustering of heatmaps was done in Morpheus.

**Figure 2- figure supplement 5. Expression profiles of TIGIT in blood and tumour and cell frequency validation with scRNA seq.** (A) Median expression of TIGIT and PD-1 on different CD8 populations in the tumour and blood. (B) Median expression of TIGIT and ICOS on different Treg populations in the tumour and blood. (C) UMAP of all cell clusters in scRNA dataset before extracting the T-cell cluster for further analysis in Fig. 3. (D) Cell frequencies from scRNA dataset. (E) PD-L1 median expression on immune, epithelial and stroma cells.

**Figure 4- figure supplement 1. Multiplex cell annotation scheme.** (A) Cancer region showing H&E (left), fluorescence image (middle) and cell annotation from HALO (right). Cyan- epithelium; magenta- αSMA, green- $CD4^+$, orange- $CD8^+$, yellow- $CD3^+$, purple- $Foxp3^+$, blue- $DAPI^+$. (B) Distribution of T-cells across stroma regions with different αSMA densities, where high was defined as the intensity from a blood vessel ($CD8^+$ T-cells, left; $CD4^+$ T-cells middle and Tregs- right).

A

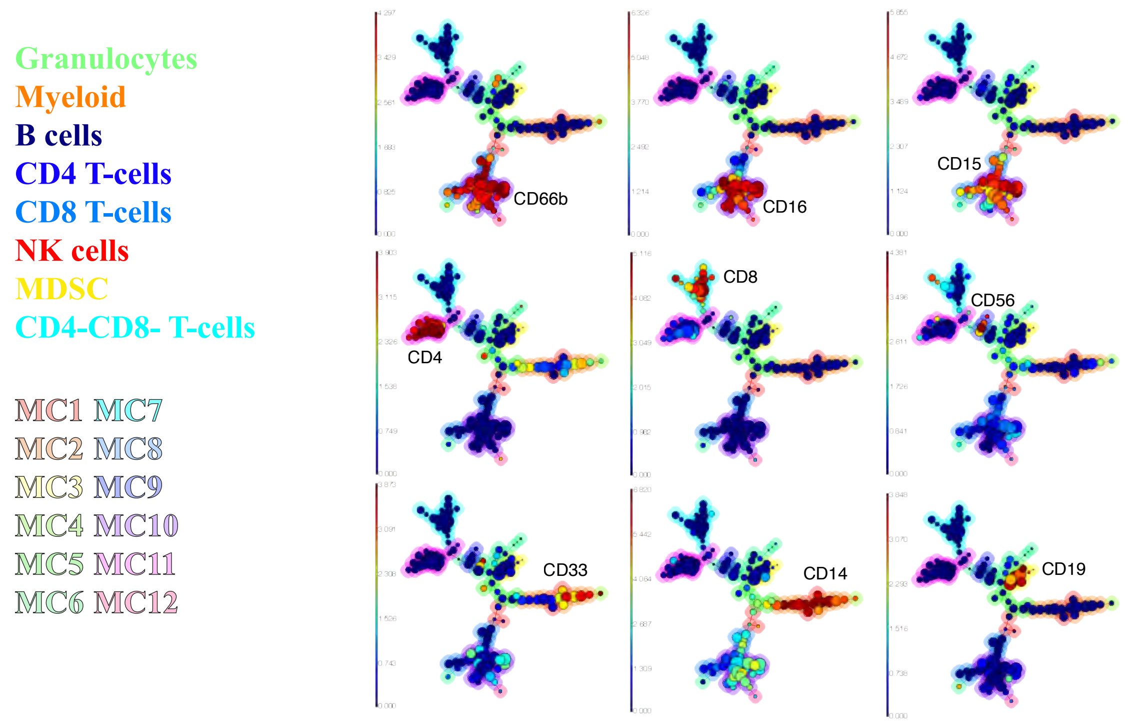


B

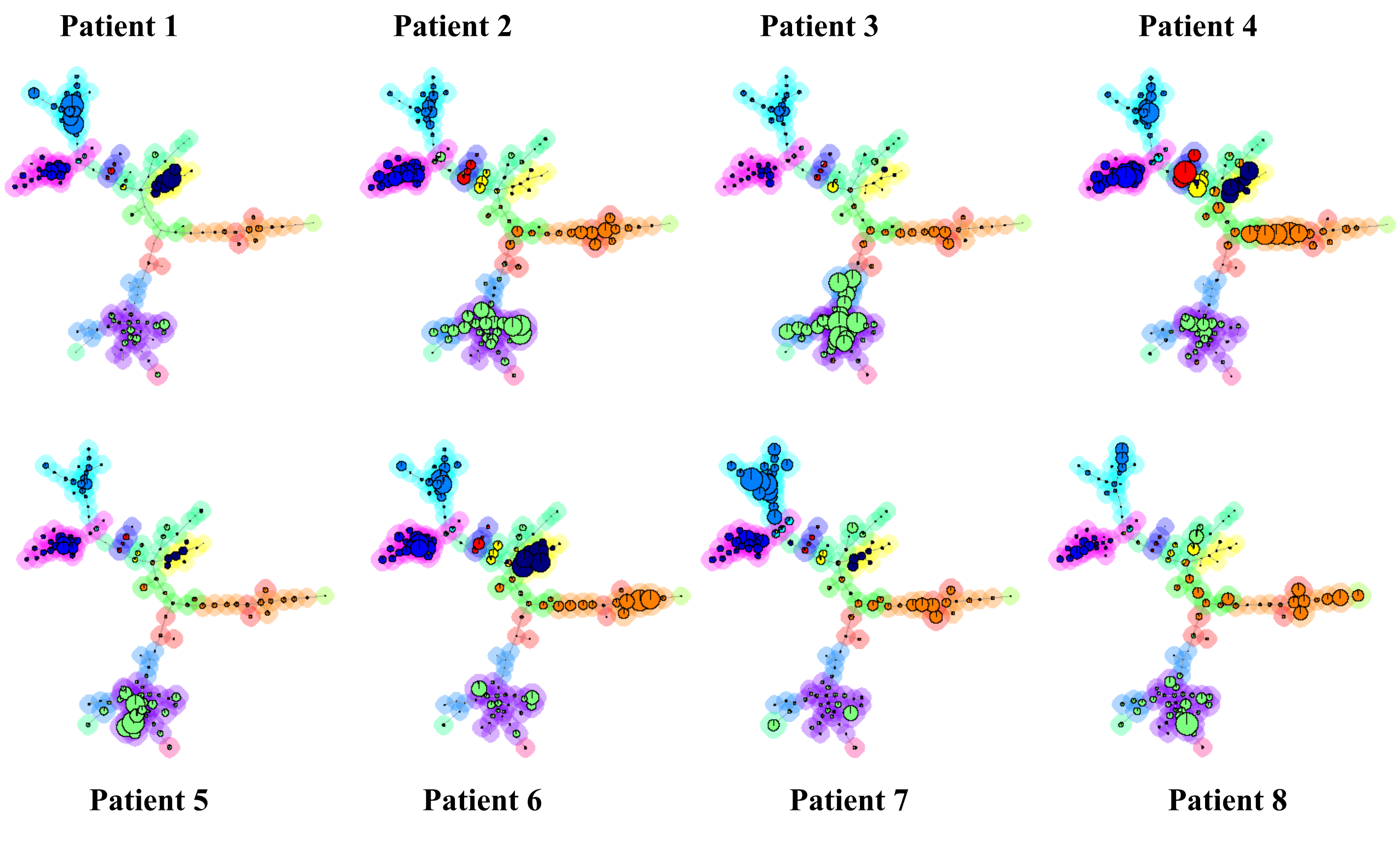


Figure S1. Sivakumar and Abu-Shah et al

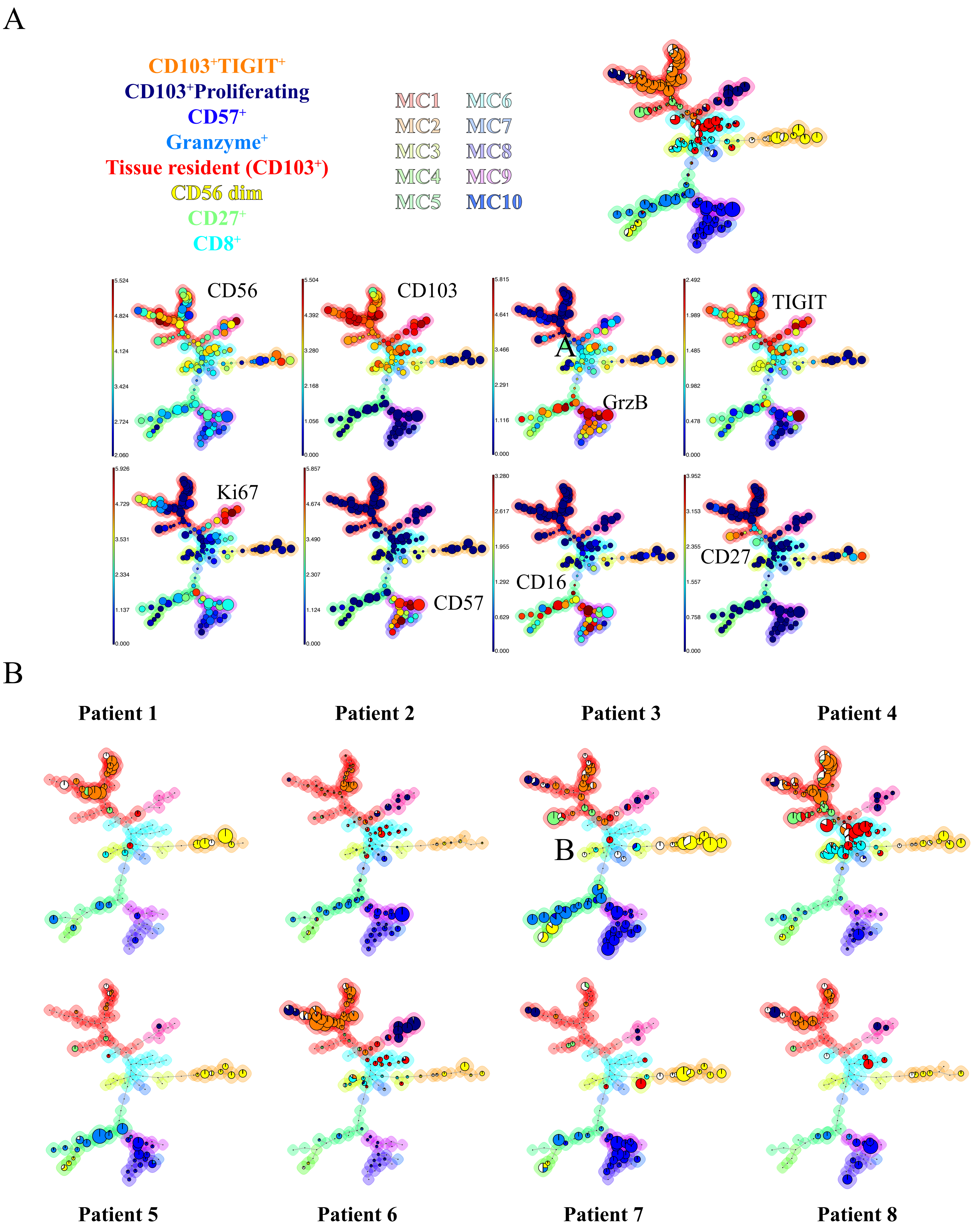


**Figure S2. Sivakumar and Abu-Shah et al**

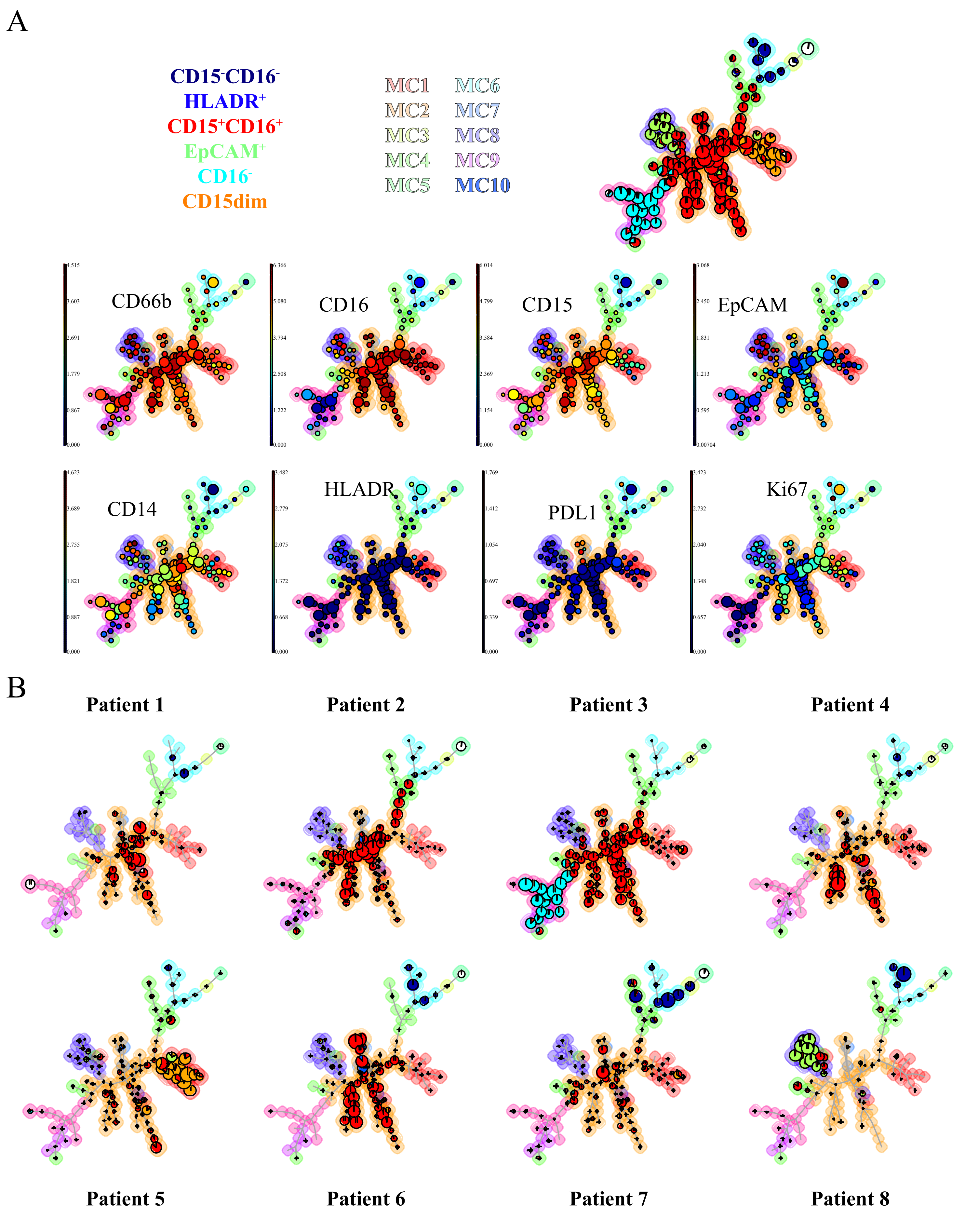


**Figure S3. Sivakumar and Abu-Shah et al**

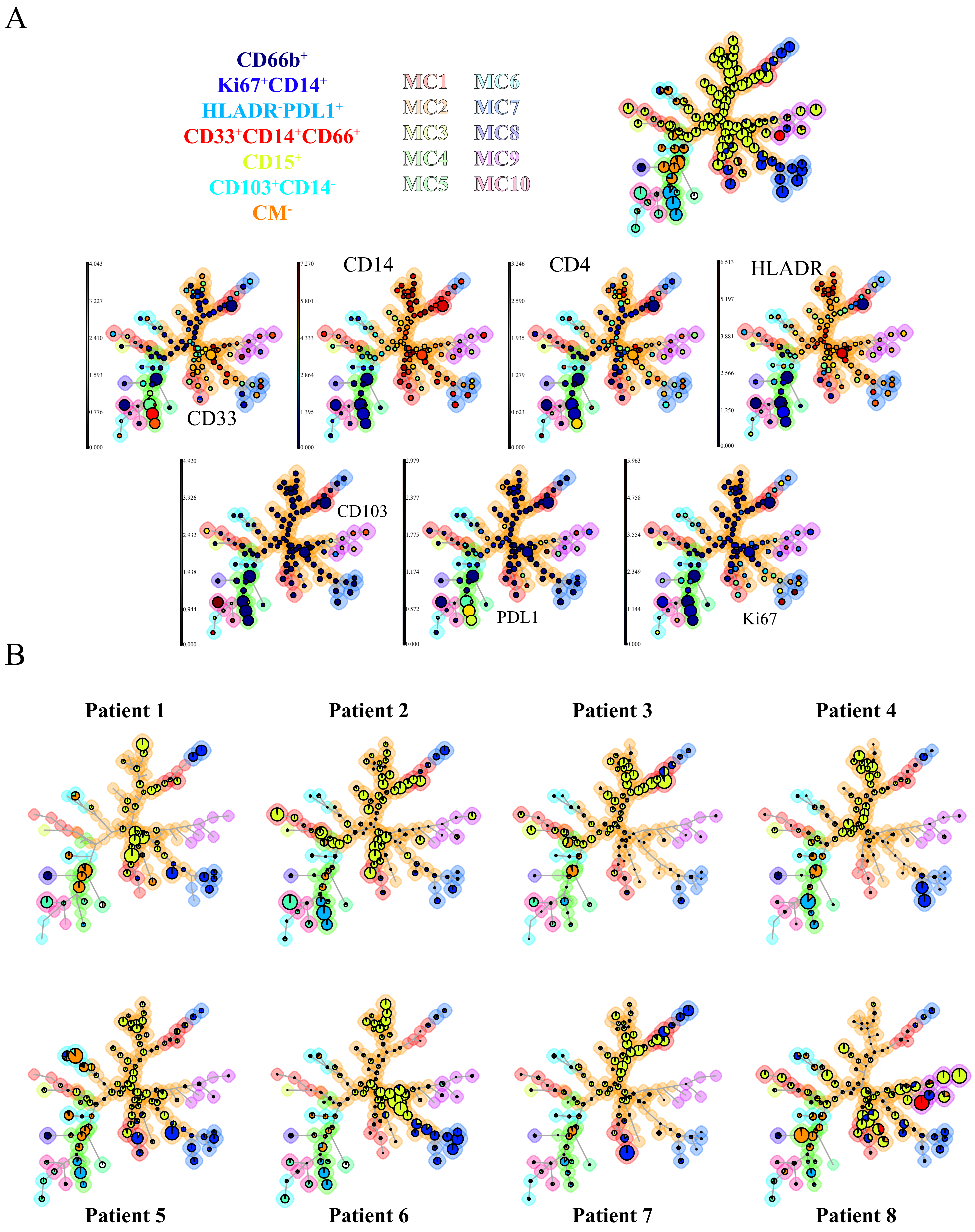


**Figure S4. Sivakumar and Abu-Shah et al**

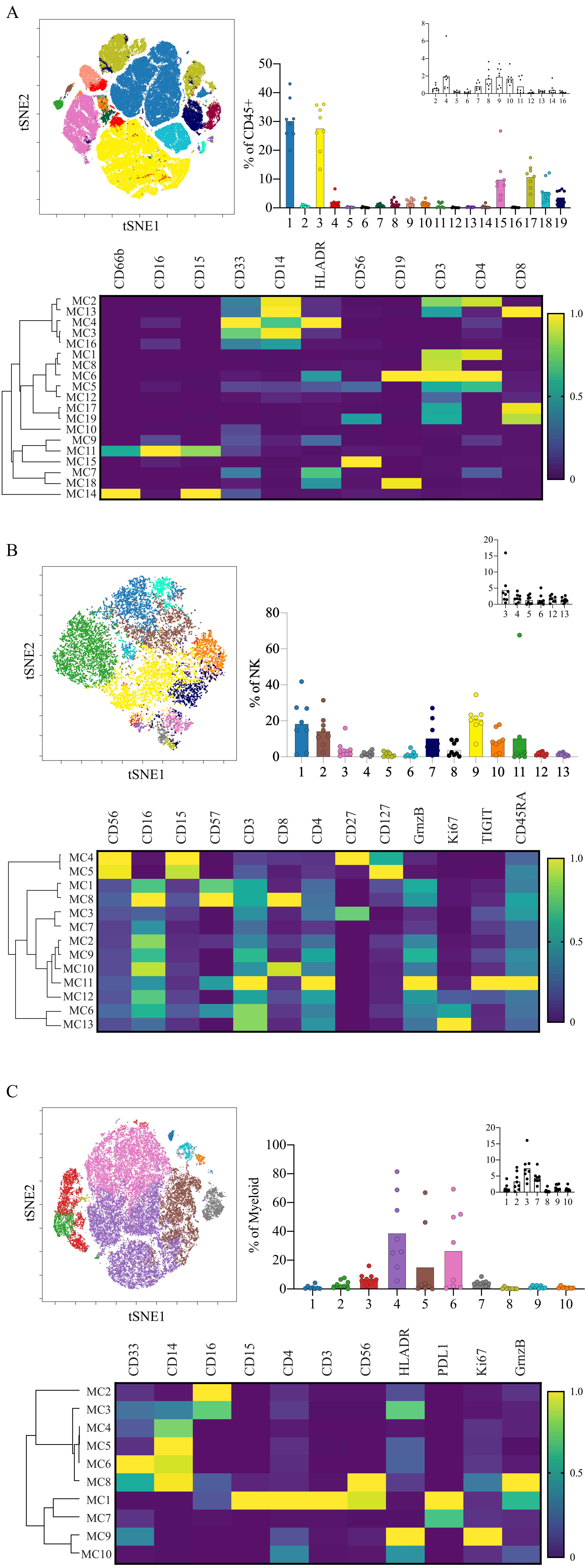


Figure S5. Sivakumar and Abu-Shah et al

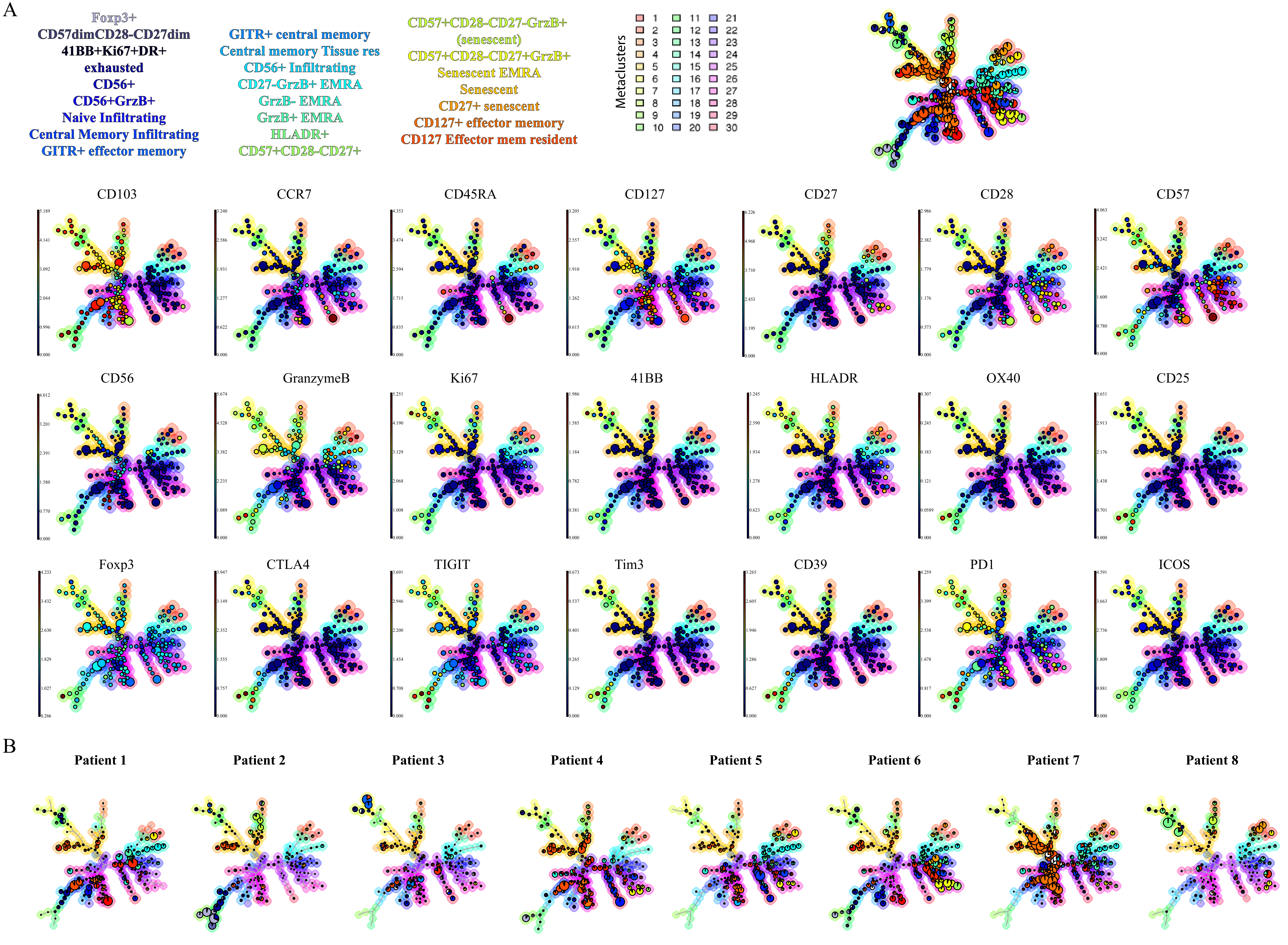


Figure S6. Sivakumar and Abu-Shah et al

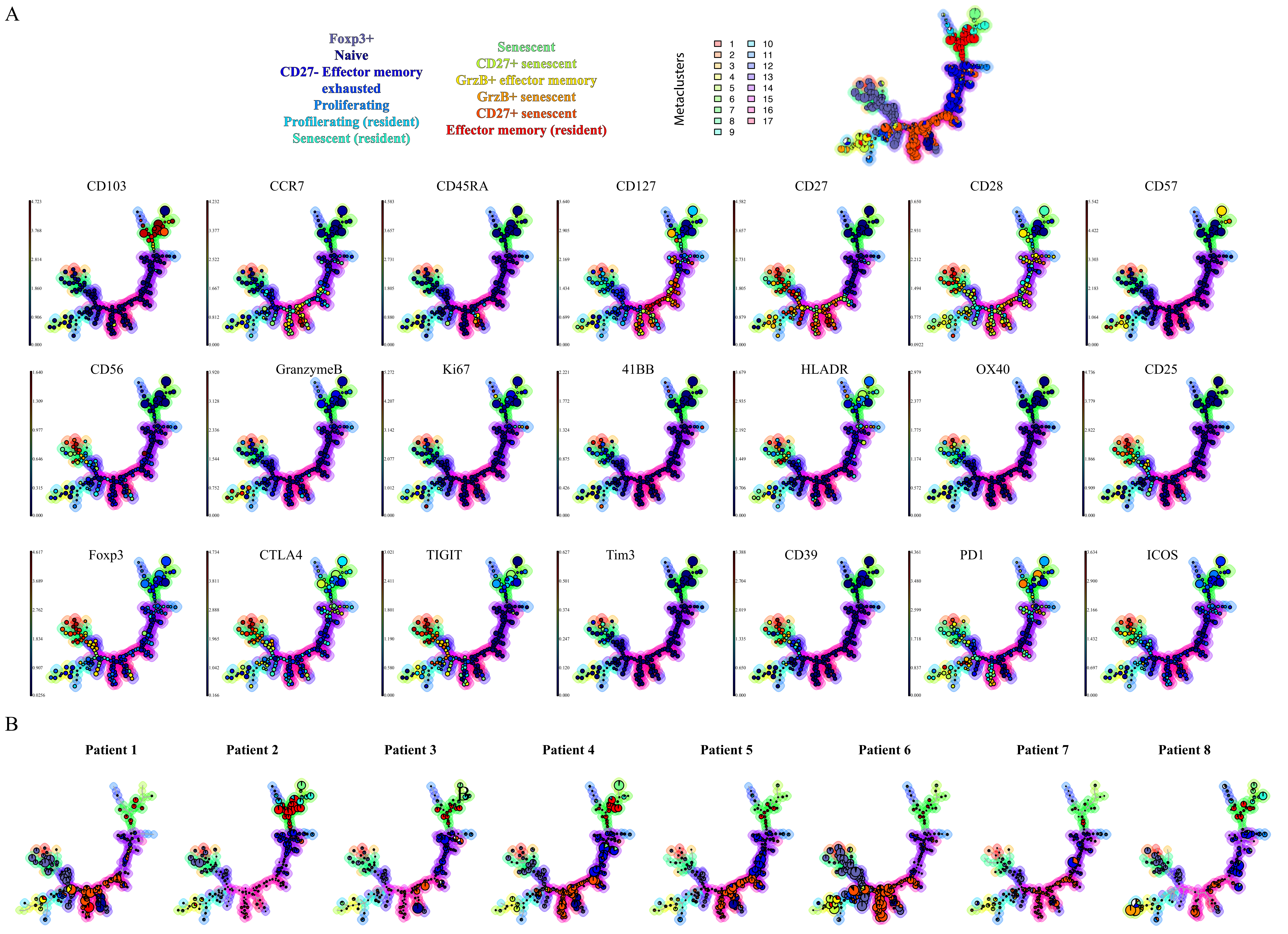


Figure S7. Sivakumar and Abu-Shah et al

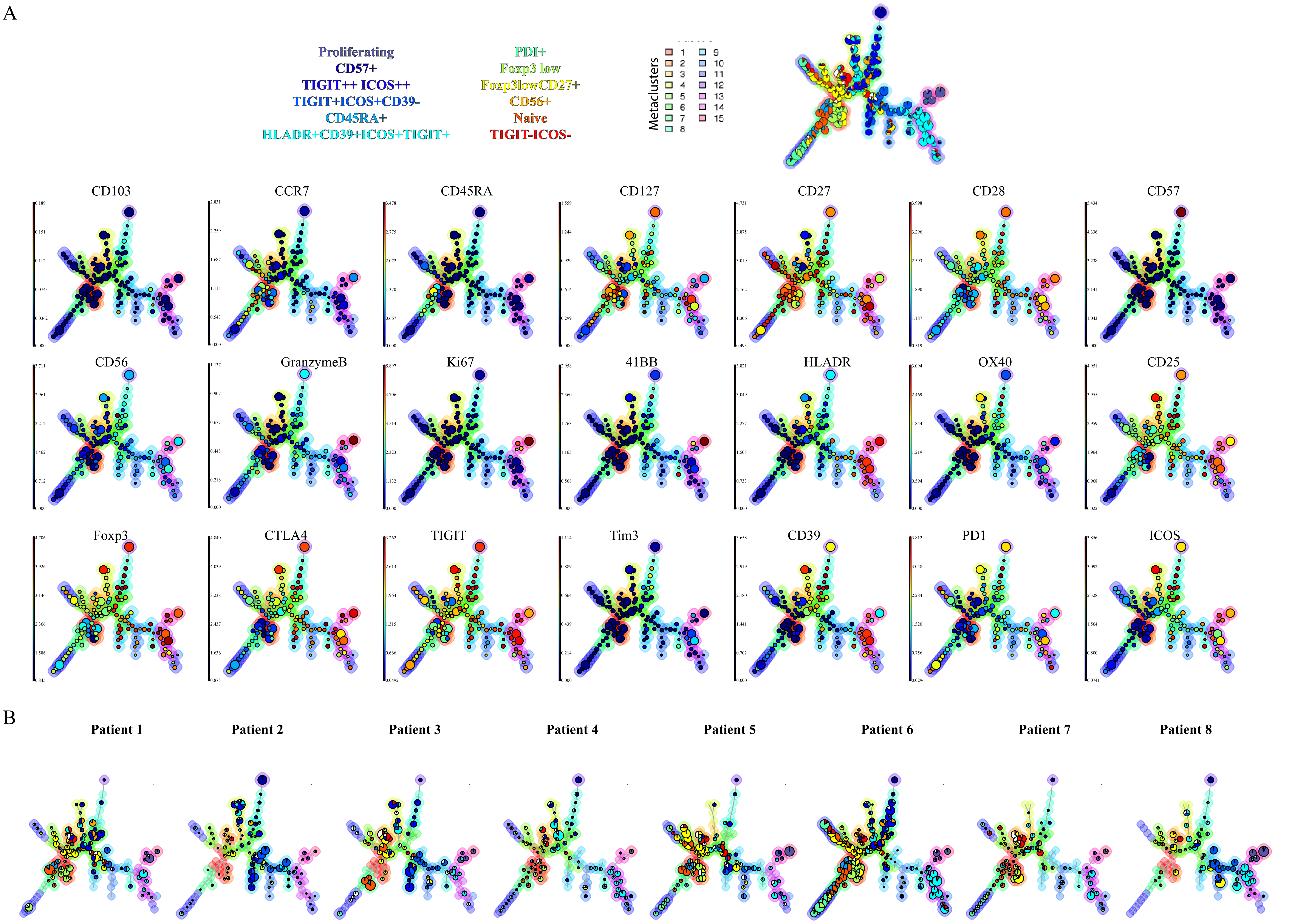


Figure S8. Sivakumar and Abu-Shah et al

**Figure S8. Individual patients' Treg populations.** Expression of clustering markers on top of a FlowSOM tree showing the relationship between metacluster, for the pooled single-cell data (top). Population frequency is presented in the pie chart in each circle along the tree, the background colour represents the metacluster identity. *MC1 (in Fig. 2C)= MC14+15 (in S8); MC3= MC1+2+5; MC6=MC3+4+6+13; MC7=MC9+10.* (Bottom) FlowSOM trees showing the individual metacluster frequency for each patient. The size of the pie chart corresponds to the frequency relative frequency. (Expression profile: from low (Dark blue) to high (Deep red)).

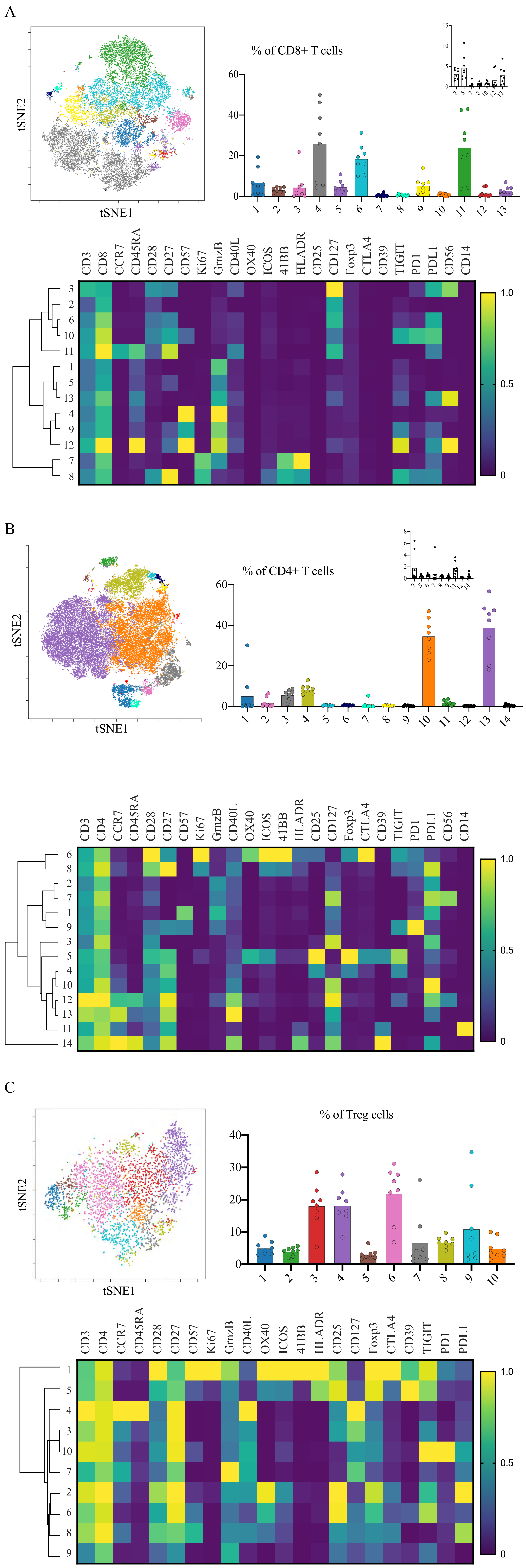


**Figure S9, Sivakumar and Abu-Shah et al**

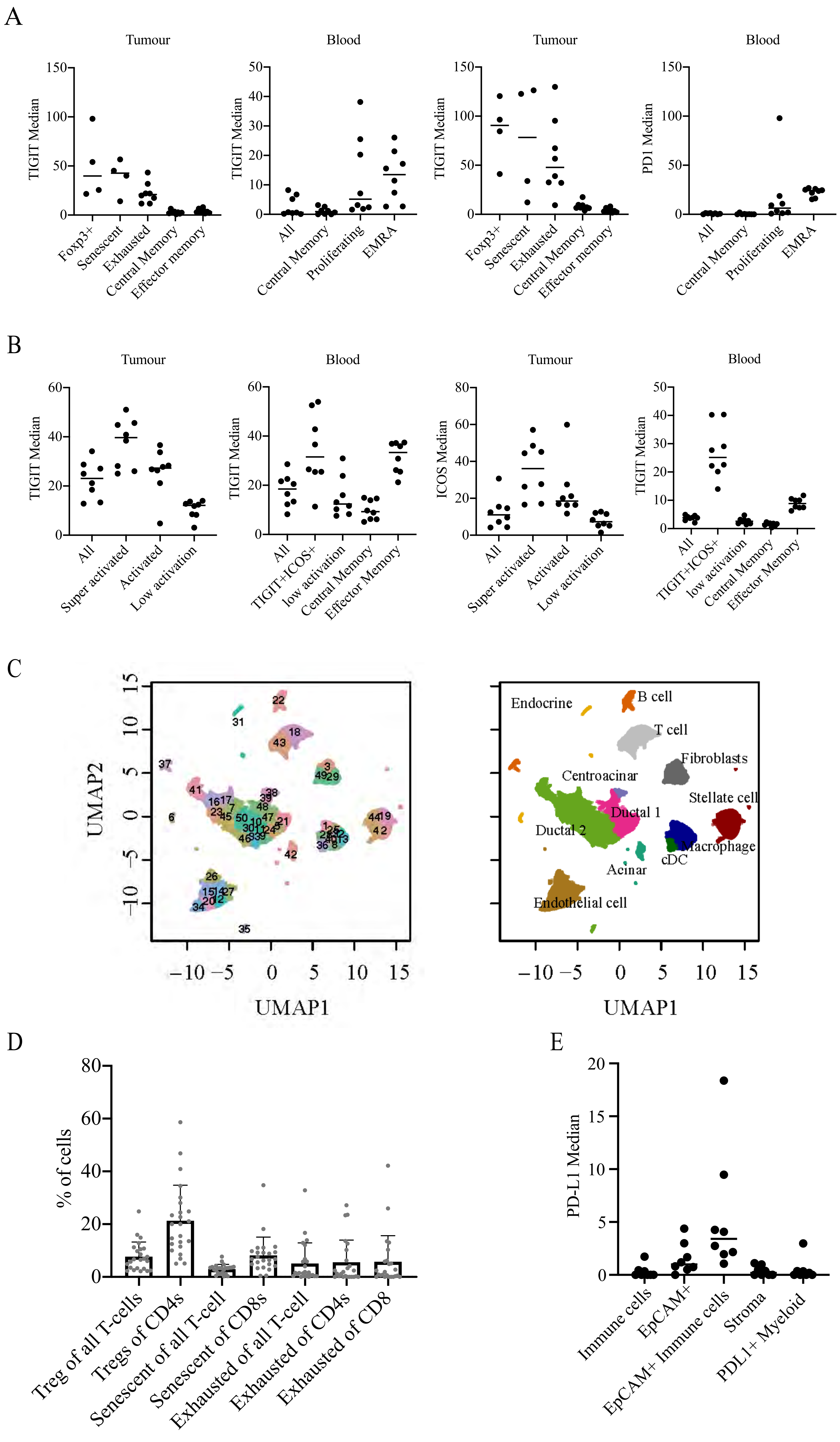


Figure S10, Sivakumar and Abu-Shah et al

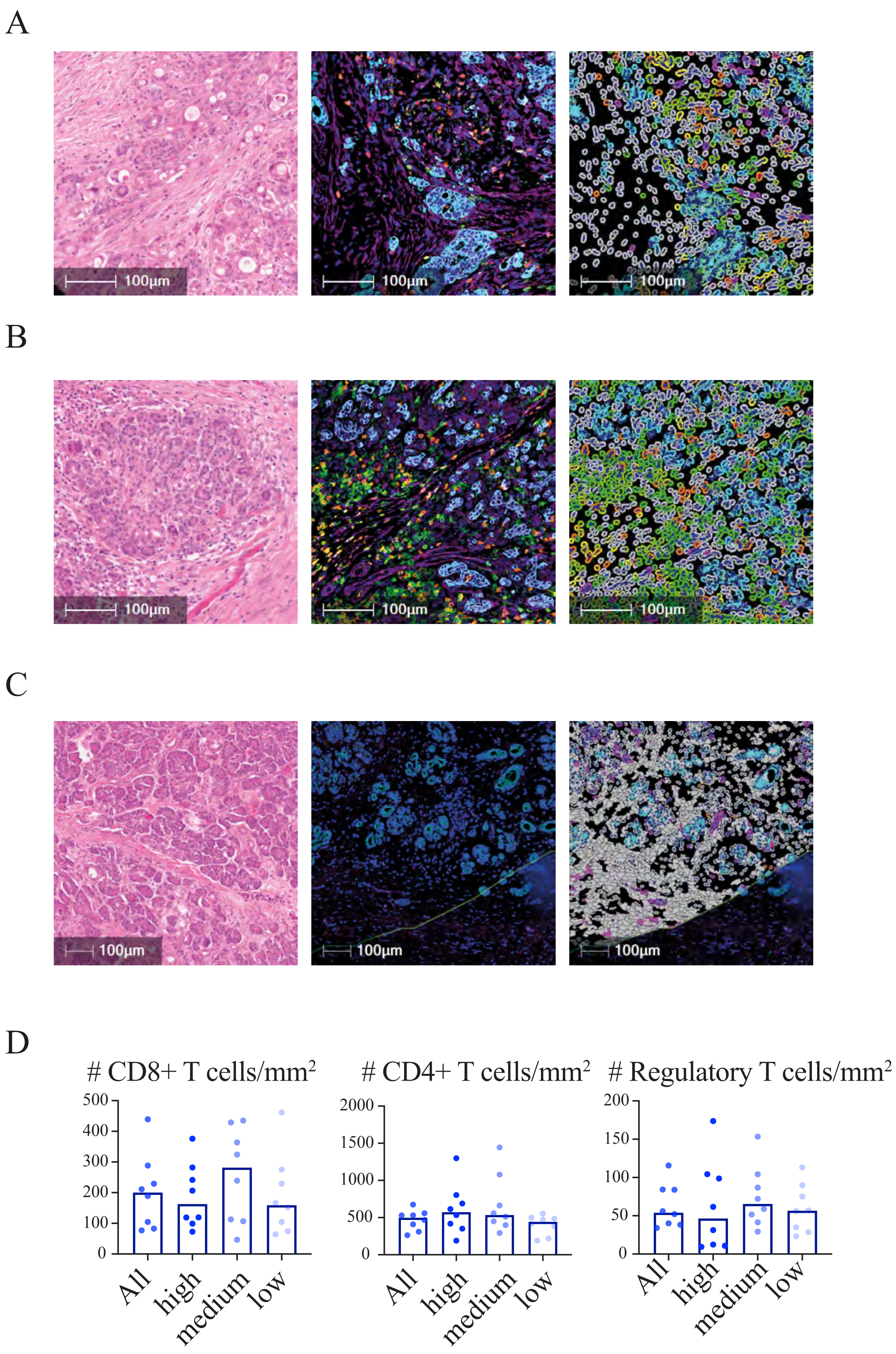


**Figure S11, Sivakumar and Abu-Shah et al**
